# On electrical spiking of *Ganoderma resinaceum*

**DOI:** 10.1101/2021.06.18.449000

**Authors:** Andrew Adamatzky, Antoni Gandia

**Author notes:** At the time of experiments AG was affiliated with Mogu S.r.l., Inarzo, Italy.

## Abstract

Fungi exhibit action-potential like spiking activity. Up to date most electrical activity of oyster fungi has been characterised in sufficient detail. It remains unclear if there are any patterns of electrical activity specific only for a certain set of species or if all fungi share the same ‘language’ of electrical signalling. We use pairs of differential electrodes to record extracellular electrical activity of the antler-like sporocarps of the polypore fungus *Ganoderma resinaceum*. The patterns of the electrical activity are analysed in terms of frequency of spiking and parameters of the spikes. The indicators of the propagation of electrical activity are also highlighted.

## 1. Introduction

Action-potential spikes are an essential component of the information processing system in a nervous system [1, 2, 3, 4, 5]. Creatures without a nervous system — plants, slime moulds and fungi — might also use the spikes of electrical potential for coordination and decision-making. There is mounting evidence that plants use the electrical spikes for a long-distance communication aimed to coordinate the activity of their bodies [6, 7, 8]. The spikes of electrical potential in plants relate to a motor activity [9, 10, 11, 12], responses to changes in temperature [13], osmotic environment [14] and mechanical stimulation [15, 16]. There is evidence of electrical current participation in the interactions between mycelium and plant roots during formation of mycorrhiza [17].

Oscillations of electrical potential in slime mould *Physarum polycephalum* were discovered in the late 1940s [18] and studied extensively [19, 20, 21]. It was demonstrated that patterns of electrical activity of the slime mould change in response to stimulation with volatile chemicals, wavelength of light, and tactile stimulation [22, 23, 24, 25, 26].

In 1976 Slayman, Long and Gradmann discovered action potential-like spikes using intra-cellular recording of mycelium of *Neurospora crassa* [27]. *Periods of* spikes evidenced in [27] were 0.2–2 min^2^. In experiments with recording of electrical potential of oyster fungi *Pleurotus djamor* we discovered two types of spiking activity: high-frequency (period 2.6 min) and low-freq (period 14 min) [29]. In semi-automated analysis of the electrical spiking activity of the hemp substrate colonised by mycelium of *Pleurotus djamor* we found that a predominant spike width is c. 6 min [30].

A question arises — “Are characteristics and patterns of electrical potential spiking the same for all species of fungi or there are some species specific parameters of the fungal electrical activity?”. Aiming to find an answer we recorded and analysed electrical activity of *Ganoderma resinaceum* sporocarps. We have been studying the electrical activity of this fungus previously [31] however only electrical responses to stimulation have been analysed and endogenous electrical activity of the mycelium has been ignored.

The paper is structured as follows. Experimental setup for recording of electrical activity of fungi is introduced in Sect. 2. Statistics and geometry of electrical potential spikes are analysed in Sect. 3. The results are discussed in Sect. 4.

## 2. Methods

The *Ganoderma resinaceum* culture used in this experiment was obtained from a wild basidiocarp found on the shores of *Lago di Varese*, Lombardy (Italy) in 2018, and maintained in alternate PDA and MEA slants at MOGU S.r.l. for the last 3 years at 4 °C under the collection code 019-18. For practical purposes, the aforementioned *G. resinaceum* culture was propagated on an sterile mixed substrate of hemp shives and soybean hulls (3:1), with a moisture content (MC) of 65% contained in plastic filter-patch microboxes c. 17 × 17 *cm*^2^ (SacO2, Belgium). The substrates were incubated in darkness at ambient room temperature c. 22°C, until antler-shaped sporocarps started to form on the surface c. 14 days post inoculation.

Electrical activity of a forming antler-like sporocarp was recorded using pairs of iridium-coated stainless steel sub-dermal needle electrodes (Spes Medica S.r.l., Italy), with twisted cables and ADC-24 (Pico Technology, UK) high-resolution data logger with a 24-bit A/D converter, galvanic isolation and software-selectable sample rates all contribute to a superior noise-free resolution. Each pair of electrodes (*i, i*+1), called ‘channel’, reported a potential difference *p*_*i*+1_*-p*_*i*_ between the electrodes. The pairs of electrodes were pierced into the sporocarp as shown in Fig. 3. The channels were from the top of the antler-like sporocarp to the bottom. Distance between electrodes was 1-2 cm. In each trial, we recorded 8 electrode pairs simultaneously. We recorded electrical activity one sample per second. During the recording, the logger has been doing as many measurements as possible (typically up to 600 per second) and saving the average value. The acquisition voltage range was 78 mV.

## 3. Results

Distribution of spike widths versus amplitudes is shown in Fig. 2(a) with the majority of spikes having amplitude less than 4 mV and width less than 41 min. Distributions of frequencies of occurrences of the spike amplitudes and widths are shown in Figs. 2(b) and 2(c). Most common amplitudes lie between 0.1 mV and 0.4 mV. The frequency distribution of the amplitudes is well described by a power 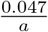, where *a* is an amplitude, shown by a solid line in Fig. 2(b). Most common widths of spikes are 300–500 sec (5–8 mins) as evidenced in Fig. 2(c). Most common types of nontrivial electrical activity observed are singular spikes, compound spikes, and trains of spikes. Singular spikes are shown by arrows in Fig. 3(a). Compound spikes are thought to be composed of several singular spikes, or a spike train which duration is in the range of a single spike width, so several spikes overlap, as labelled by arrows with stars in Fig. 3(a). An example of the train of spikes is shown in Fig. 3(a). In the first train the average distance between spikes is 731 sec and in the second train 2300 sec. There are two potential origins of the compound spikes. First, the compound spikes could be seen as trains of spikes with a very short distance between the spikes, i.e. trains of overlapping spikes. Second, the compound spikes could be see as a slow wave potential superimposed with a train of low amplitude spikes. Single spikes can form analogies of damped or growing type oscillators. The growing type of oscillator with increasing amplitude and period of oscillations is illustrated by four spikes in Fig. 3(c). The amplitude increases as 0.35 mV, 0.84 mV, 1.01 mV, 1.27 mV. The width of spikes increases as 88 sec, 210 sec, 252 sec, 273 sec. The distance between spikes is changing as 2253 sec, 3739 sec, 5372 sec.

**Figure 1:**
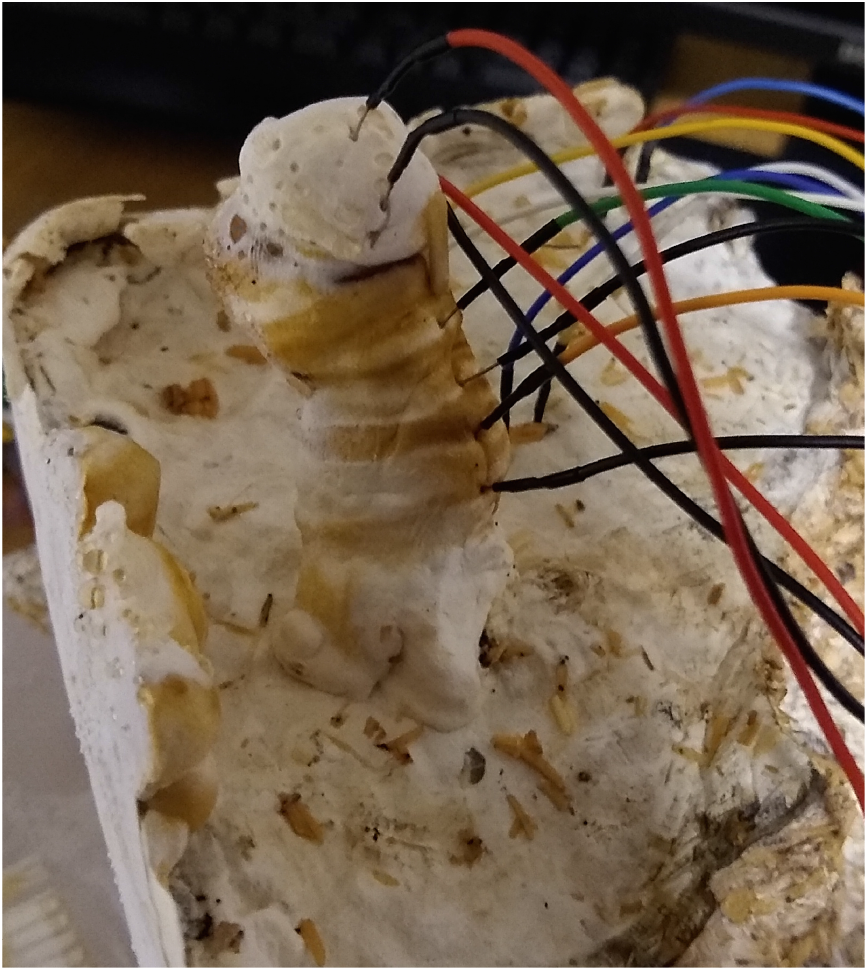
Position of electrodes in a sporocarp of *G. resinaceum*.

**Figure 2:**
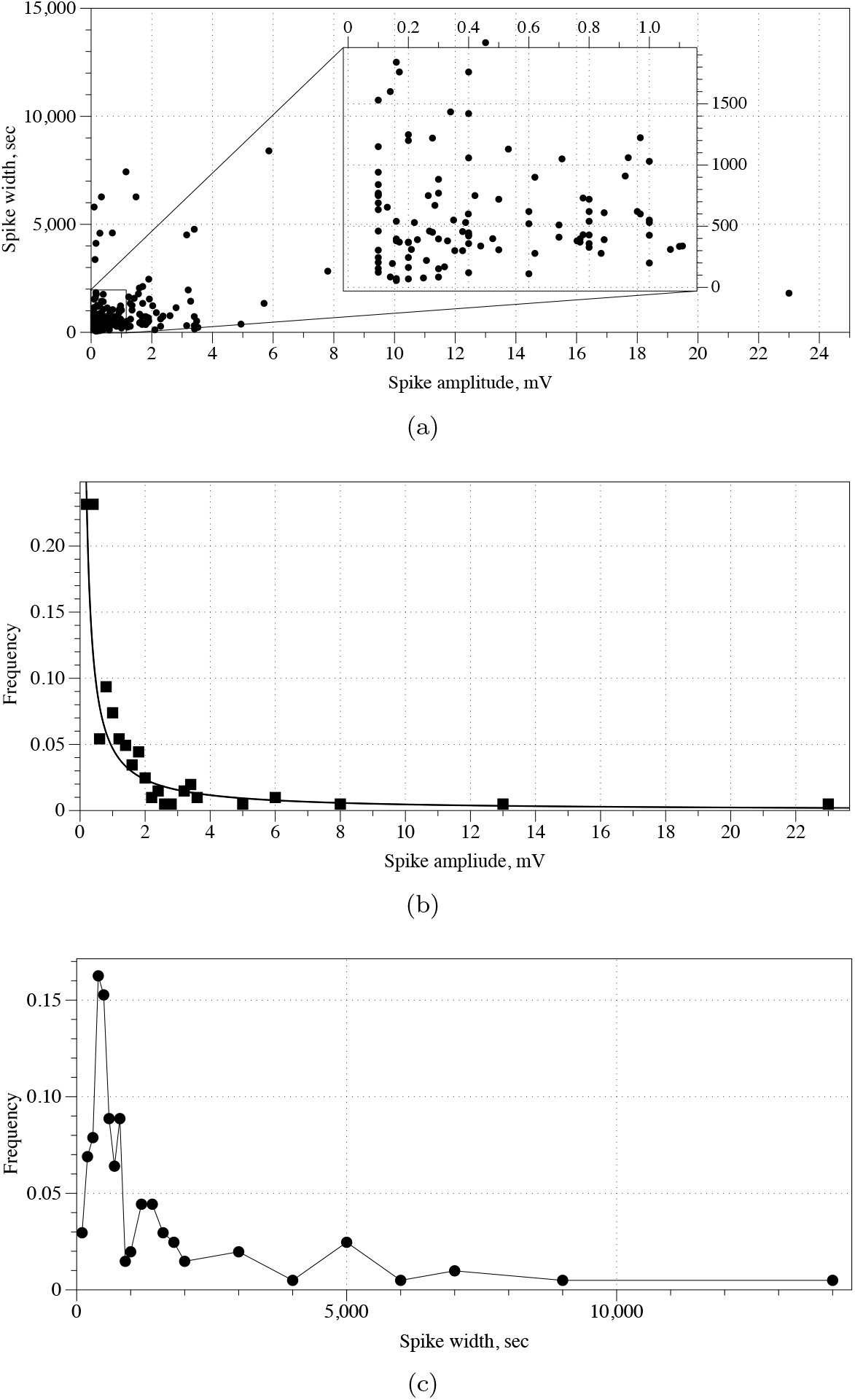
Quantitative analysis of *G. resinaceum* spiking activity. (a) Amplitude versus width for three sample channels, recorded for 60 hours. Distribution of spike (b) amplitudes and (c) widths. Distribution of amplitudes is approximated by a power law 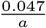, where *a* is an amplitude, shown by curve in (b)

**Figure 3:**
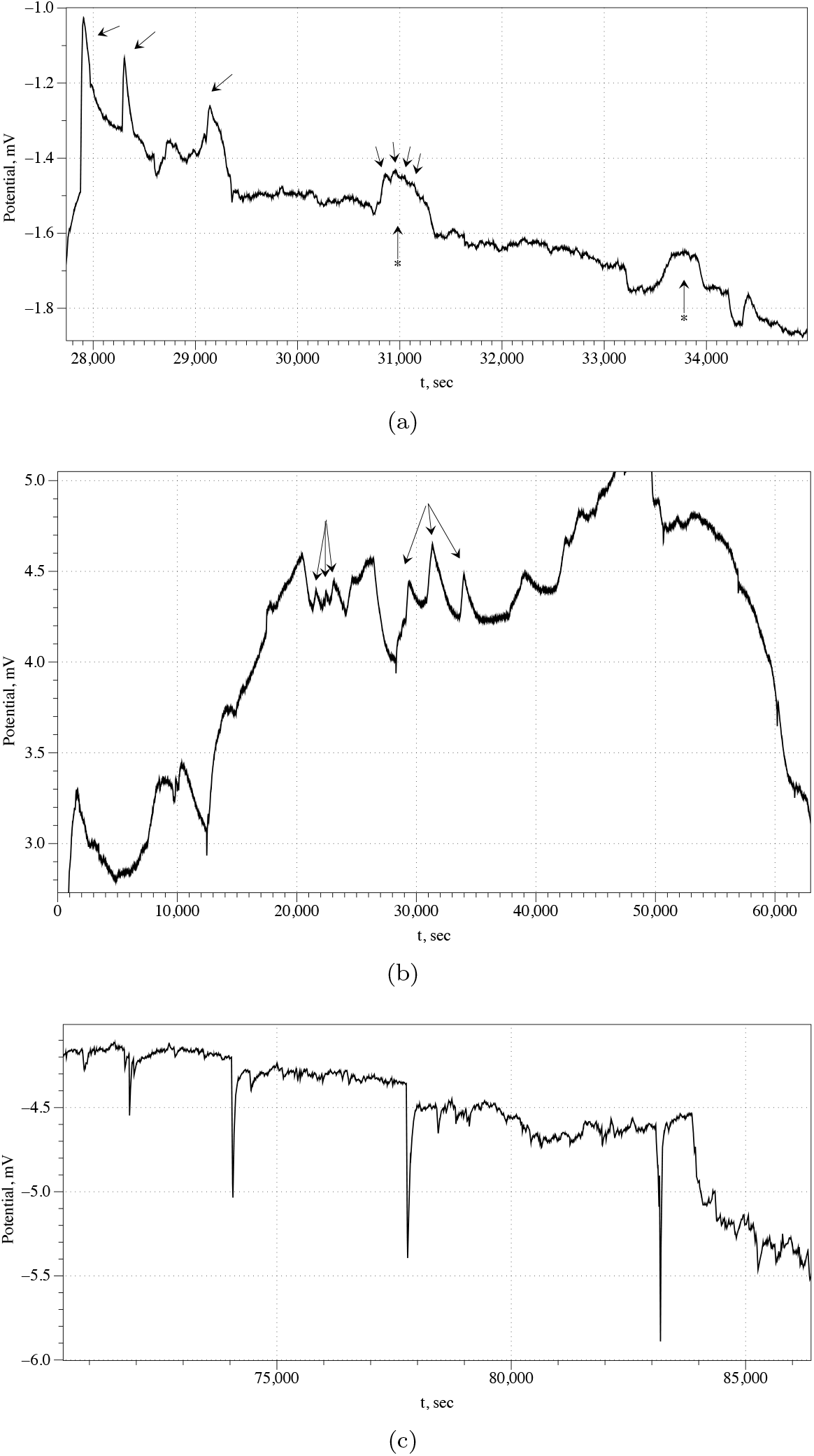
Examples of an electrical activity of *G. resinaceum* sporocarp. (a) Single and compound spikes. Compound spike is shown by arrow with start. (b) Examples of trains of spikes. Two trains of three spikes each are shown by arrow. (c) Growing type oscillator.

A rare but interesting feature observed is a long (up to 2 hours) burst of high-frequency (a spike per 7 min) spikes. An example is shown in Fig. 4. The main burst is preceded by a short (13 min) burst of five spikes, shown as ‘S’ in Fig. 4, and is completed by another short (15 min) burst of seven spikes. There are c. 70 spikes in the main burst: average width of a spike is 29 sec and average amplitude 0.1 mV; an average distance spikes is 92 sec.

**Figure 4:**
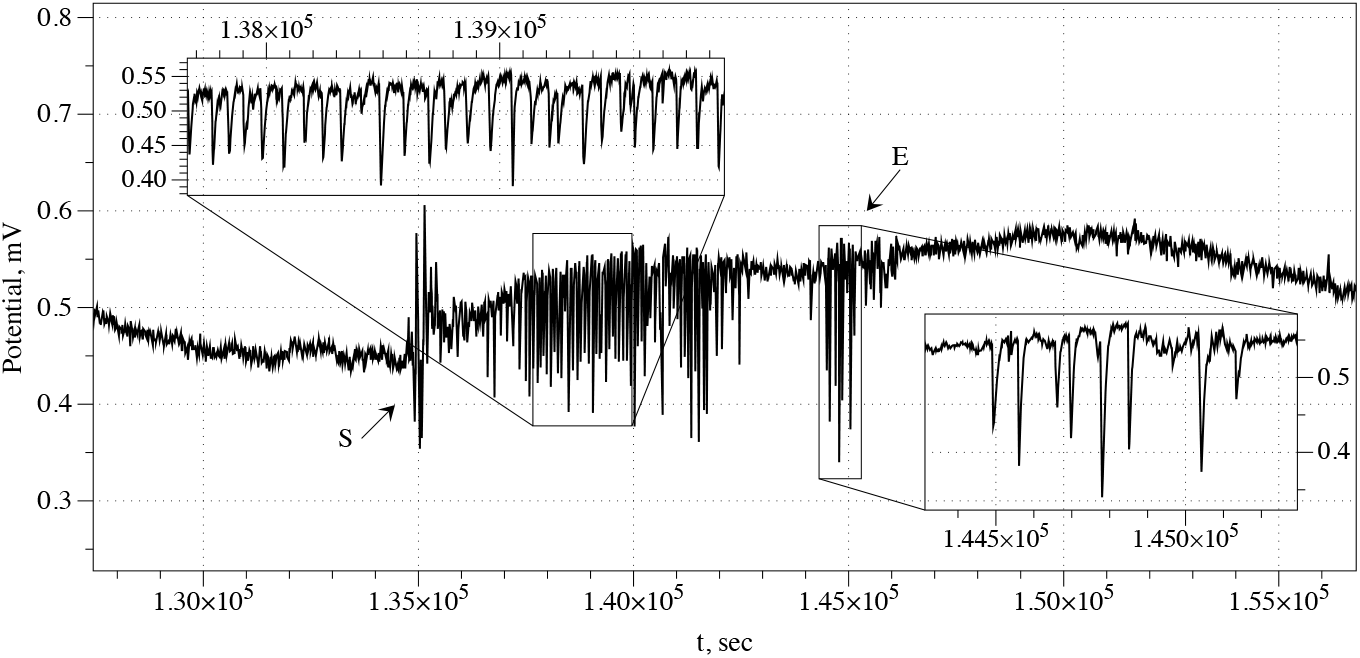
Example of a long high frequency burst of spikes.

## 4. Discussion

We found that most common width of an electrical potential spike is 5-8 min. An average distance between electrodes in the pairs of differential electrodes was 1 cm. We speculate it takes a phase-wave, which governs the electrical potential, to propagate between the electrodes 6 min in average, i.e. a speed of the phase-wave is c. 0.028 mm/sec. Calcium waves can propagate with the fastest speed up 0.03 mm/s [32]. Thus we can speculate that the spikes of the electrical potential in *Ganoderma resinaceum* antler-like sporocarps might relate to the fast calcium waves. The nature of long burst of high-frequency (a spike per 7 mins) short-width (half-a-minute) spikes is beyond speculations and could not be attributed to calcium waves.

The width of the spike of antler-like sporocarps of *Ganoderma resinaceum* is twice of the width of the spikes detected in sporocarps of *Pleurotus djamor* [29] *yet of the same width as spikes detected in a substrate colonised by mycelium of Pleurotus djamor* [30]. *Possible explanations are the following. First, sporocarps show higher growth rate than mycelium. Therefore, spikes detected on sporocarps of Pleurotus djamor* are twice as narrow as spikes detected on a substrate colonised by mycelium of *Pleurotus djamor*. Second, *Ganoderma resinaceum* might have a slower metabolism than *Pleurotus djamor* and therefore its spiking activity shows wider spikes, and consequently, lower frequency of the spiking.

Recalling our initial question — “Do different species of fungi have different parameters of their electrical spiking activity?” — we could answer “Likely, yes”. However it would be reckless to base the answer on comparing just two species of fungi. Therefore future research will focus on collecting statistics of spiking of wider range of fungi species.

## Acknowledgement

This project has received funding from the European Union’s Horizon 2020 research and innovation programme FET OPEN “Challenging current thinking” under grant agreement No 858132. The authors would like to acknowledge the collaboration of Mogu S.r.l. providing the living materials used in the experiments.

## Footnotes

2 In 1995 Olsson and Hansson demonstrated spontaneous action potential like activity in a hypha of *Pleurotus ostreatus* and *Armillaria bulbosa* (synonymous with *A. gallica* and *A. lutea*) via intra-cellular recording with a reference electrode in an agar substrate [28]. Our present results concern extracellular recordings, therefore we will not compare thee with Olsson and Hansson results.

